# Adaptive 2.5D base-pairing subgraph search detects RNA small-molecule binding sites

**DOI:** 10.64898/2026.07.07.737024

**Authors:** David Nitchi, Jérôme Waldispühl, Carlos Oliver

## Abstract

`Ribo-LENS` is a geometric deep-learning framework for detecting small-molecule binding sites in RNA structures. It is designed to exploit two properties of RNA base-pairing networks: their robustness to conformational fluctuation and the functional signatures they encode. By reasoning directly in the space of base-pairing subgraphs, `Ribo-LENS` assembles coherent binding sites, in contrast to methods that score residues independently. In extensive experiments, `Ribo-LENS` is competitive with, and often outperforms, large all-atom co-folding models (AlphaFold3, Chai-1), fine-tuned language models (GerNA-Bind), and structure-based tools (RNAsite), raising mean MCC to 0.380 (versus 0.321 for the state-of-the-art GerNA-Bind). It is strongly robust to apo/holo rearrangement, with its accuracy tracking the base-pairing graph (Spearman *ρ* = 0.82 with binding-site graph edit distance) rather than backbone displacement (*ρ* = −0.15 with RMSD), and depends far less on sequence homology than competing predictors. In an end-to-end, sequence-based virtual screen of the ROBIN assay (∼25,000 compounds), `Ribo-LENS` guides docking to a small predicted pocket, matching a blind all-atom cavity search (enrichment factors up to 6.1) at a fraction of the search cost; on two SARS-CoV-2 targets its predicted sites align with NMR chemical-shift perturbations. `Ribo-LENS` turns coarse base-pairing structure into a practical entry point for screening the vast, largely unexplored RNA target space.

## Introduction

Ribonucleic acids (RNAs) are an under-exploited reservoir of therapeutic targets. The majority of the human genome is transcribed, but only a small fraction is translated into protein. To date, large-scale transcriptome surveys estimate that on the order of 75–80% of the genome gives rise to RNA, while protein-coding exons account for less than 2%^(1)^. Still, most drugs target proteins and the set of clinically validated, druggable protein targets covers only a few percent of the proteome^(2)^. Against this backdrop, if even a fraction of functional non-coding RNAs proves druggable, the pool of targets amenable to structure-based drug discovery would exceed its protein counterpart by orders of magnitude. However, currently only a handful of RNA targets are being actively pursued in drug-discovery campaigns, among which we find bacterial riboswitches, viral structured elements such as the HIV-1 TAR and trans-activation response elements, and the SARS-CoV-2 frameshift pseudoknot^(3)^. This gap is at least partially attributable to the difficulty of accurately identifying target sites, and the development of efficient binding-site detection methods for traversing the transcriptome is therefore a prerequisite for unlocking RNA as a target class.

Binding-site detection in RNA is uniquely challenging. Rule-based heuristics established for proteins translate poorly: protein pockets occupy surface cavities, whereas RNA binding sites are more often buried within the folded architecture and coincide with locally flexible regions that confer binding specificity^(4)^. The chemical space of RNA binders is also distinct, more positively charged and reliant on hydrogen-bonding and stacking, so ligand-based, protein-calibrated approaches transfer poorly ^(5;3)^.

A more fundamental obstacle is the scarcity of high-resolution 3D structures, a prerequisite for docking and for training the machine-learning models needed to screen libraries that now exceed a billion compounds ^(6)^. Even when structures exist, many are apo (ligand-free) forms ill-suited to docking because binding often induces rearrangement, and modest modeling errors in a holo form can derail prediction.

To address this, we previously proposed abstracting RNA structure as its base-pairing network. These interactions, classified geometrically by Leontis and Westhof^(7)^, give a robust signature of the 3D fold while being far less sensitive to conformational fluctuation than all-atom representations. We call this network, comprising the Watson–Crick, wobble and non-canonical base pairs together with backbone connectivity, the RNA’s *2*.*5D graph* ^(8;9)^, and showed it encodes signal to score RNA–small-molecule complexes and accelerate virtual screening by orders of magnitude^(10)^. Those methods, however, presuppose a known binding site, unavailable for most transcripts. Base pairs are also faster to predict than full 3D structures^(11)^, enabling high-throughput pipelines.

Existing binding-site predictors fall short on RNA in characteristic ways. Sequence-based methods exploit a signal that is too weak for reliable RNA pocket localization; structure-based methods either depend on hand-crafted features or score residues independently, and so cannot search the combinatorial space of connected substructures in which binding sites actually reside. In principle, binding sites could be discovered directly from sequence via all-atom co-folding models such as AlphaFold3 ^(12)^ and Chai-1 ^(13)^, but co-folding of RNA–ligand complexes has yet to demonstrate consistent success.

In this paper we introduce `Ribo-LENS`, a 2.5D graph-based method for predicting the small-molecule binding sites of an RNA. Unlike residue-level predictors, which assign each nucleotide an independent binding probability, `Ribo-LENS` constructs each site as a single coherent unit: a learned graph-contraction procedure^(14)^ repeatedly merges high-scoring interactions into connected supernodes, growing each pocket outward from a high-confidence core until the local signal is exhausted. Because contraction terminates and can restart elsewhere, `Ribo-LENS` naturally predicts *multiple* distinct sites per molecule; and by scoring whole substructures rather than isolated residues, it reasons directly in the combinatorial space of connected substructures in which binding sites actually reside, capturing whether a residue raises the binding likelihood in the context of a particular group.

Although it uses only a coarse-grained, base-pairing representation of RNA structure, `Ribo-LENS` outperforms both specialized RNA predictors (GerNA-Bind, RNAsite) and all-atom co-folding methods (AlphaFold3, Chai-1) on a held-out structural benchmark^(15)^, and its accuracy depends far less on sequence homology to the training set. On paired apo/holo structures, its predictions track changes in the binding-site *graph* rather than 3D displacement, remaining stable even under large (*>* 10 Å) backbone rearrangements. In an end-to-end, fully sequence-based screening setting, coupling `Ribo-LENS` with predicted structures and pocket-restricted docking matches blind docking’s enrichment against an experimental assay of ∼25,000 compounds ^(5)^ while confining the search to a small, size-independent region, and on two therapeutically relevant SARS-CoV-2 elements `Ribo-LENS` localizes binding sites that agree with experimentally reported ligand-perturbed residues. Notably, although `Ribo-LENS` is trained only on RNA–ligand co-crystal structures from the PDB, the binding signal it recovers is reflected across three independent sources of evidence: structure-based docking, the *in vitro* ROBIN small-molecule binding assay, and NMR chemical-shift perturbations. This suggests it captures genuine binding determinants that generalize beyond its crystallographic training data. Combined with our previous contributions ^(10)^, this work moves toward the massive RNA virtual-screening applications needed to realize the potential of RNAs as pharmaceutical targets.

## Results

### Ribo-LENS learns to construct coherent binding sites from base-pairing networks

`Ribo-LENS` is a geometric deep-learning framework that identifies small-molecule binding sites directly from RNA structure (Fig. 2). Each RNA is encoded as a relational graph in which nodes are nucleotides and edges are base-pairing interactions typed by the Leontis-Westhof classification^(7)^, augmented with backbone connectivity. This 2.5D representation captures the interaction geometry that defines binding pockets while remaining far less sensitive than all-atom models to the conformational fluctuations that complicate docking. Each nucleotide additionally carries a one-hot identity encoding and an evolutionary embedding from the RNA-FM language model^(16)^. To localize binding sites, we couple Relational Graph Convolutional Network (RGCN) layers ^(17)^ with an adaptive pooling module derived from EdgePool ^(14)^. The RGCN embeds each nucleotide and assigns it a binding score; the embedding and score are concatenated and passed to an edge-scoring network that ranks each interaction by its proximity to a binding site. The pooling module iteratively contracts high-scoring edges, merging connected nucleotides into supernodes representing coherent pockets (Fig. 1a); training drives each contracted region to be enriched for binding-site nucleotides (Methods). This gives two advantages over node-level predictors (Fig. 1b): sites are returned as connected subgraphs rather than scattered residues, and the model scores whole *substructures*, capturing whether a residue contributes to a pocket in the context of its group.

**Figure 1:**
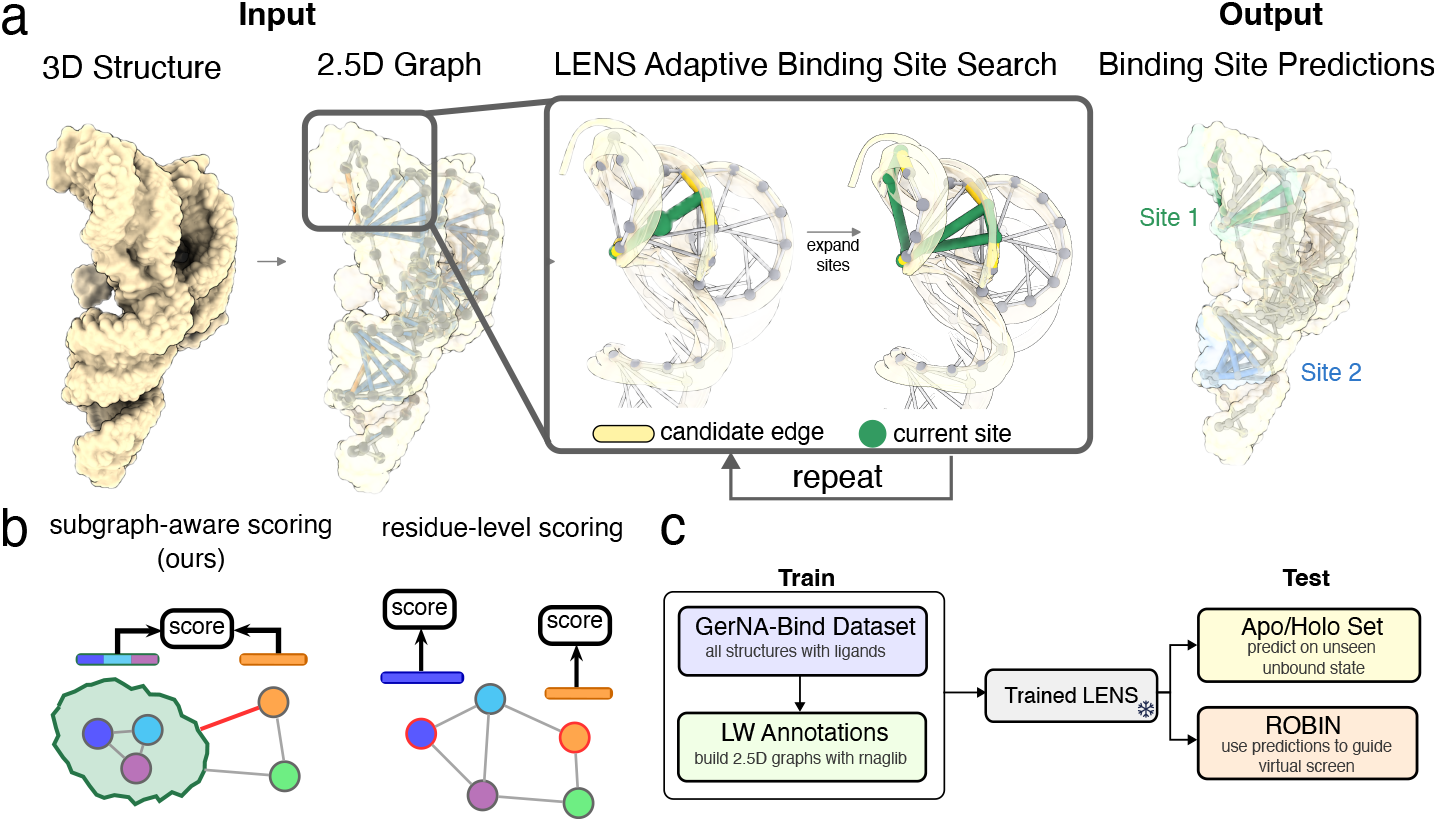
Overview of binding site prediction pipeline. (a) `Ribo-LENS` pooling procedure. (b) Conceptual difference between subgraph-level (group-aware) binding-site reasoning and independent residue-wise prediction. (c) Dataset construction: we use RNAGlib to retrieve RNA–ligand structures from the PDB, label binding residues by a distance cutoff, convert structures to 2.5D graphs, and perform redundancy-aware structure-based splitting.

**Figure 2:**
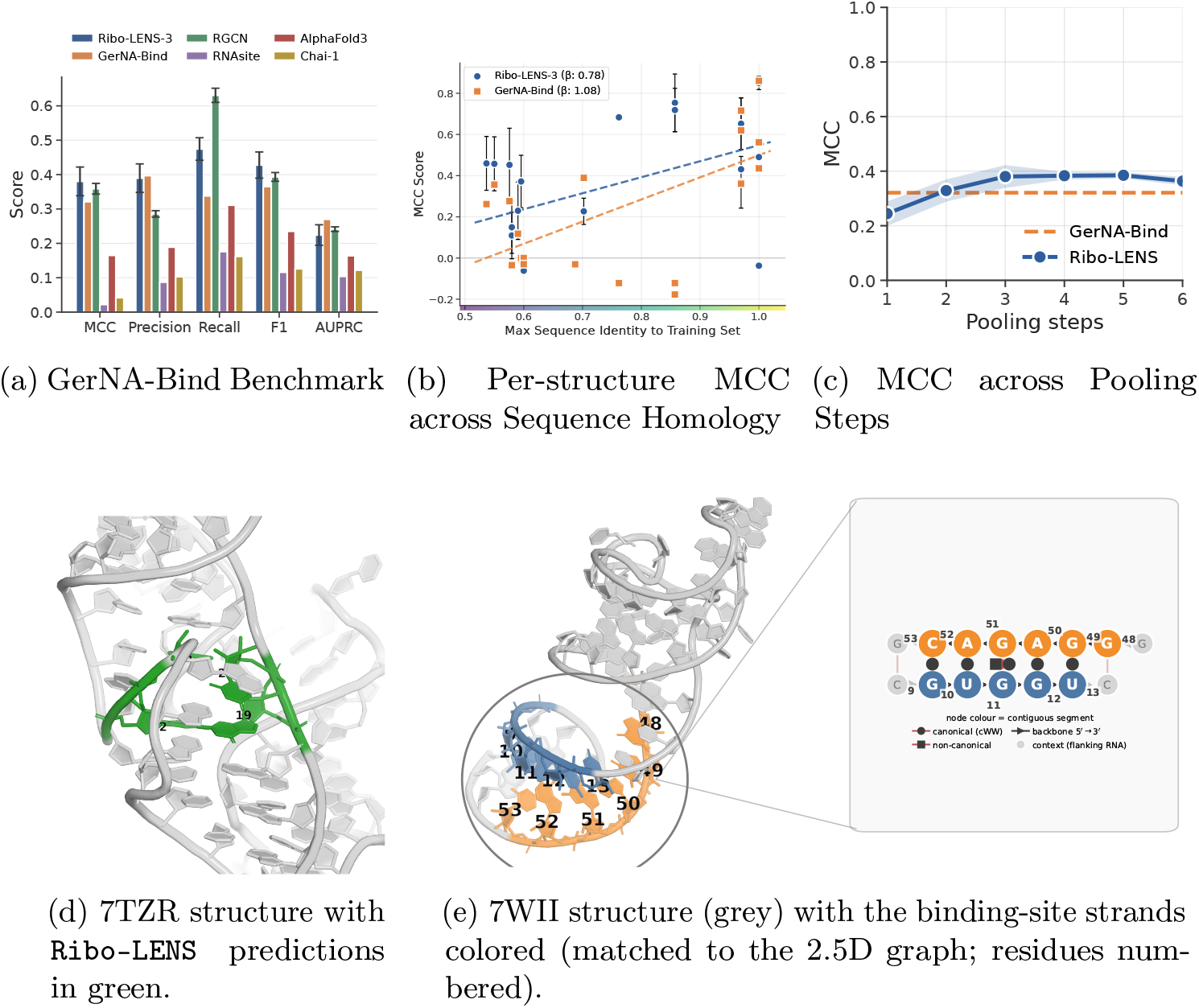
Ribo-LENS on the GerNA-Bind test set (*n* = 22 structures). **(a)** Mean MCC, precision, recall, F1 and AUPRC for `Ribo-LENS`-3 (`Ribo-LENS` at three pooling steps), GerNA-Bind, the underlying RGCN, RNAsite, AlphaFold3 and Chai-1; error bars denote the standard deviation across 9 random seeds. **(b)** Per-structure MCC versus maximum sequence identity to the training set for `Ribo-LENS`-3 (blue circles) and GerNA-Bind (orange squares), with linear-regression trend lines (`Ribo-LENS`: *β* = 0.78, *R*^2^ = 0.21, *p <* 0.001; GerNA-Bind: *β* = 1.08, *R*^2^ = 0.39, *p* = 0.002). **(c)** MCC on the test set as a function of pooling steps for `Ribo-LENS` (blue line; shaded band, standard deviation across 9 seeds) and GerNA-Bind (orange dashed line); the corresponding precision and recall sweeps are shown in Fig. 2. **(d)** Structure of 7TZR with the four nucleotides predicted by `Ribo-LENS` in green; all lie within 3 Å of the ligand yet none overlap the two annotated binding residues, illustrating a sparse-labeling artifact rather than a mis-localization. **(e)** The 7WII RNA (grey cartoon) with the two contiguous strands of the `Ribo-LENS`-predicted binding site colored (matched to its 2.5D graph in the magnified panel; residues numbered). The motif is an internal loop built from an A–G *cis* Watson–Crick and a G–G *cis* Watson–Crick/Hoogsteen pair.

### Ribo-LENS outperforms specialized and general-purpose structure-based predictors

We first benchmark `Ribo-LENS` against state-of-the-art predictors, then examine how performance depends on training-set homology and on the learnable graph-contraction architecture.

#### Benchmark comparison

Figure 2a evaluates `Ribo-LENS` on the GerNA-Bind test set. We compare `Ribo-LENS` at three pooling steps (matching training; hereafter `Ribo-LENS`-3) and its underlying RGCN against two specialized RNA predictors (GerNA-Bind, RNAsite) and two all-atom co-folding models (AlphaFold3, Chai-1), which receive the RNA sequence and the ligand SMILES and jointly predict the complex.

Despite operating on only a coarse-grained representation, both `Ribo-LENS`-3 and the RGCN outperform RNAsite, AlphaFold3 and Chai-1 on every metric, showing that a coarse-grained base-pairing representation is sufficient to surpass current co-folding models on this task. `Ribo-LENS`-3 attains the highest mean MCC (0.380, versus 0.359 for the RGCN and 0.321 for GerNA-Bind ^(15)^) and the highest F1 score. The comparison to the bare RGCN isolates the effect of graph contraction. Pooling refines the RGCN’s node-level predictions into more balanced pockets by trading the RGCN’s very high recall (0.63, at low precision 0.285) for a more balanced operating point (precision 0.390, recall 0.474), yielding higher MCC overall. The advantage over GerNA-Bind is driven by recall: the two reach comparable precision (0.390 vs. 0.397) while `Ribo-LENS` recovers far more binding residues (0.474 vs. 0.338). The only metric on which `Ribo-LENS` trails *both* GerNA-Bind and the RGCN is AUPRC, which reflects its discrete output rather than weaker localization: nucleotides outside a predicted pocket all score zero, so any missed binding residue cannot be recovered at any threshold, truncating the precision–recall curve even where the predicted pockets are accurate.

#### Ribo-LENS’s accuracy is less coupled to training-set homology than GerNA-Bind’s

Figure 2b plots per-RNA MCC against each structure’s maximum sequence identity to the training set. GerNA-Bind’s accuracy rises more steeply with homology than `Ribo-LENS`’s (*β* = 1.08 vs. 0.78) and is more tightly coupled to it (*R*^2^ = 0.39 vs. 0.21): `Ribo-LENS` wins on the low- and mid-homology structures (10 of 15, one tie) while GerNA-Bind regains the advantage at high homology (4 of 7). `Ribo-LENS` is also more robust overall, returning a non-positive MCC on only 5 of 22 structures versus 10 for GerNA-Bind. Its accuracy is thus far less explained by training-set similarity, the property most relevant to deploying a predictor across the largely unannotated transcriptome. The same pattern holds at five pooling steps (Supp. Fig. 1).

All five `Ribo-LENS` failures are interpretable. Three (7WIB, 7WIF and 7WII, resolutions of the same RNA) are the only low-homology structures where GerNA-Bind wins, yet it too is negative on all three; here `Ribo-LENS` localizes an internal-loop motif (an A–G *cis* Watson–Crick adjacent to a G–G *cis* Watson– Crick/Hoogsteen pair; Fig. 2e) whose geometry is characteristic of a flexible, binding-competent site but lies away from the annotated pocket. The other two are labeling artifacts rather than mis-localizations: for 7TZR (Fig. 2d) all four predicted nucleotides lie within 3 Å of the ligand, and for 8G8Z within 4–5 Å, yet none overlap the sparsely annotated positive residues.

#### Three pooling steps balance precision and recall

Figures 2c, 2a and 2b trace MCC, precision and recall against pooling depth. MCC rises sharply over the first three steps, plateaus between steps 3 and 5 (peak 0.385 at step 5), and declines thereafter; precision tracks GerNA-Bind through the first three steps and then degrades, while recall increases monotonically. This matches the intended behavior of the mechanism, which grows each pocket outward from a high-confidence core: early steps are precise but conservative, and later steps trade precision for recall as lower-confidence nucleotides are absorbed. Three steps therefore sit near the MCC optimum while preserving precision, justifying it as the default. Across-seed variance concentrates in the first three steps and then falls (net mean-MCC standard-deviation reduction of 0.038 between steps 3 and 5; Supplementary), consistent with different initializations converging on the same pockets as contraction proceeds.

In sum, `Ribo-LENS` depends far less on training-set homology than GerNA-Bind and fails on fewer structures. Furthermore, its residual failures either surface alternative binding-competent motifs or fall within a few Å of the ligand at nucleotides the sparse reference labels omit and its predictions grow more consistent across initializations as pooling proceeds.

### Ribo-LENS is robust to apo–holo conformational change

Structure-based predictors are trained on bound (holo) states, yet at deployment we are typically given only the unbound (apo) state. Sequence-based models sidestep this by ignoring structure, but at a large cost in accuracy, as our GerNA-Bind benchmark shows. We therefore ask whether `Ribo-LENS`’s predictions survive the apo–holo conformational change when the model is provided only the apo state as input.

#### High-confidence cores translate from holo to apo

We use a hand-curated dataset^(4)^ of RNAs resolved in both states, spanning apo–holo backbone RMSDs from 0.42 to 10.72 Å. To quantify reconfiguration in graph space we compute the graph edit distance (GED) between the apo and holo binding-site graphs (Methods); because `Ribo-LENS` reasons over 2.5D graphs, which are less sensitive to rearrangement than atomic coordinates, we expect this graph-space distance, rather than backbone RMSD, to track its robustness. Across pooling steps (Fig. 3a), holo and apo MCC are nearly identical over the first few contractions and both rise monotonically, diverging only later. The divergence is a precision effect (Fig. 3b): apo and holo precision start together near 0.65 and separate after step two, while recall stays comparable (Fig. 3c). The high-confidence residues recovered early are thus essentially the same whether the model sees the bound or unbound structure; the states part only during the later, less accurate outward expansion. For a pipeline that relies chiefly on localizing a confident core, this early-stage agreement is what matters most.

**Figure 3:**
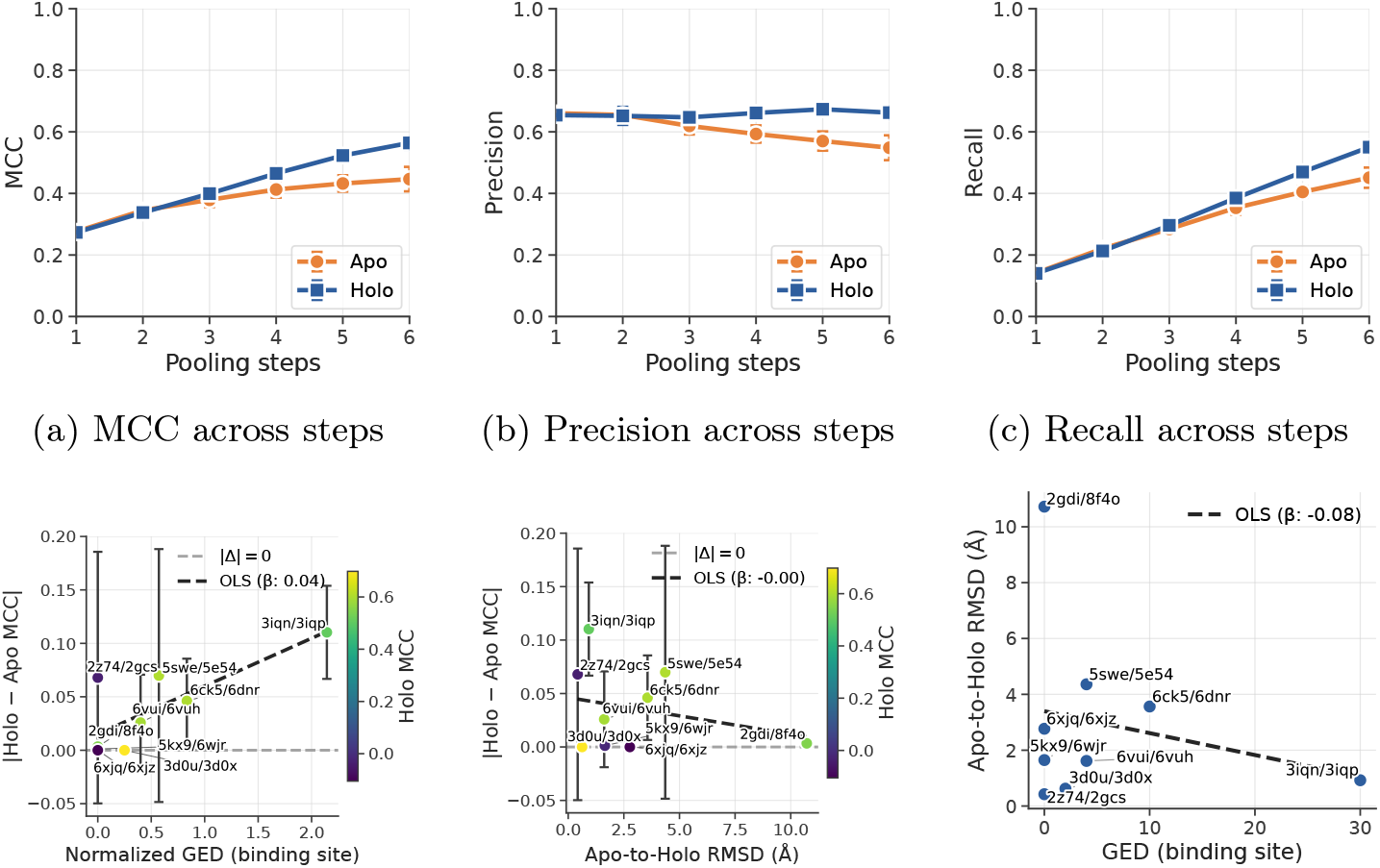
(a–c) MCC, precision and recall performance between Apo and Holo structures for `Ribo-LENS` across pooling steps. (d) Absolute delta between Holo and Apo MCC versus normalized GED of the binding-site graphs in Holo and Apo states after 3 pooling steps. Dark dashed line represents OLS trendlines (*β* = 0.047, *R*^2^ = 0.220, *p* = 9.91*×* 10^−6^; Spearman *ρ* = 0.818, *p* = 7.03 *×*10^−3^; Pearson *r* = 0.836, *p* = 5.00 *×*10^−3^). (e) Absolute delta between Holo and Apo MCC versus RMSD from the Apo to the Holo structure after 3 pooling steps. Dark dashed line represents OLS trendlines (*β* = −0.003, *R*^2^ = 0.014, *p* = 2.93 *×* 10^−1^; Spearman *ρ* = −0.150, *p* = 7.00 *×* 10^−1^; Pearson *r* = −0.211, *p* = 5.86 *×* 10^−1^).

#### Predictions transfer across states when the base-pairing network is preserved

Fixing the analysis at three pooling steps, the absolute holo–apo MCC difference is strongly associated with binding-site GED (Spearman *ρ* = 0.82, *p <* 0.01; Fig. 3d) but not with backbone RMSD (*ρ* = − 0.15, *p* = 0.70; Fig. 3e); since GED and RMSD are themselves uncorrelated (Fig. 3f; Supp. Fig. 5), the same atomic displacement need not perturb the base-pairing graph, and it is the graph perturbation that predicts `Ribo-LENS`’s change in output. The effect is specific to rewiring *within* the pocket: computing GED over a one-hop neighborhood removes it (Supp. Fig. 4). Two pairs bracket it (Fig. 4): the 2GDI/8F4O pair (Fig. 4a,4b) undergoes a large 10.72 Å rearrangement yet, with its binding-site graph conserved (GED 0.0), differs by only 0.003 in holo–apo MCC (2GDI’s holo is in the GerNA-Bind training set, but the large one-hop change means the apo prediction reflects generalization, not memorization); conversely, the 3IQN/3IQP pair (Fig. 4c,4d) differs by just 0.92 Å RMSD but, with the pocket rewired (a *trans* Hoogsteen/sugar-edge U45–A57 contact replaced by a *cis* Watson–Crick A57–U46 pair), shows the largest MCC gap (0.108) and is in fact *more* accurate on the apo form in 6 of 9 seeds. The benchmark is small (*n* = 9) and per-structure MCC is seed-variable, so we treat these associations as indicative; even so, they point to a clear principle: `Ribo-LENS` trades sensitivity to ubiquitous 3D displacement for sensitivity to the rarer event of base-pairing rewiring.

**Figure 4:**
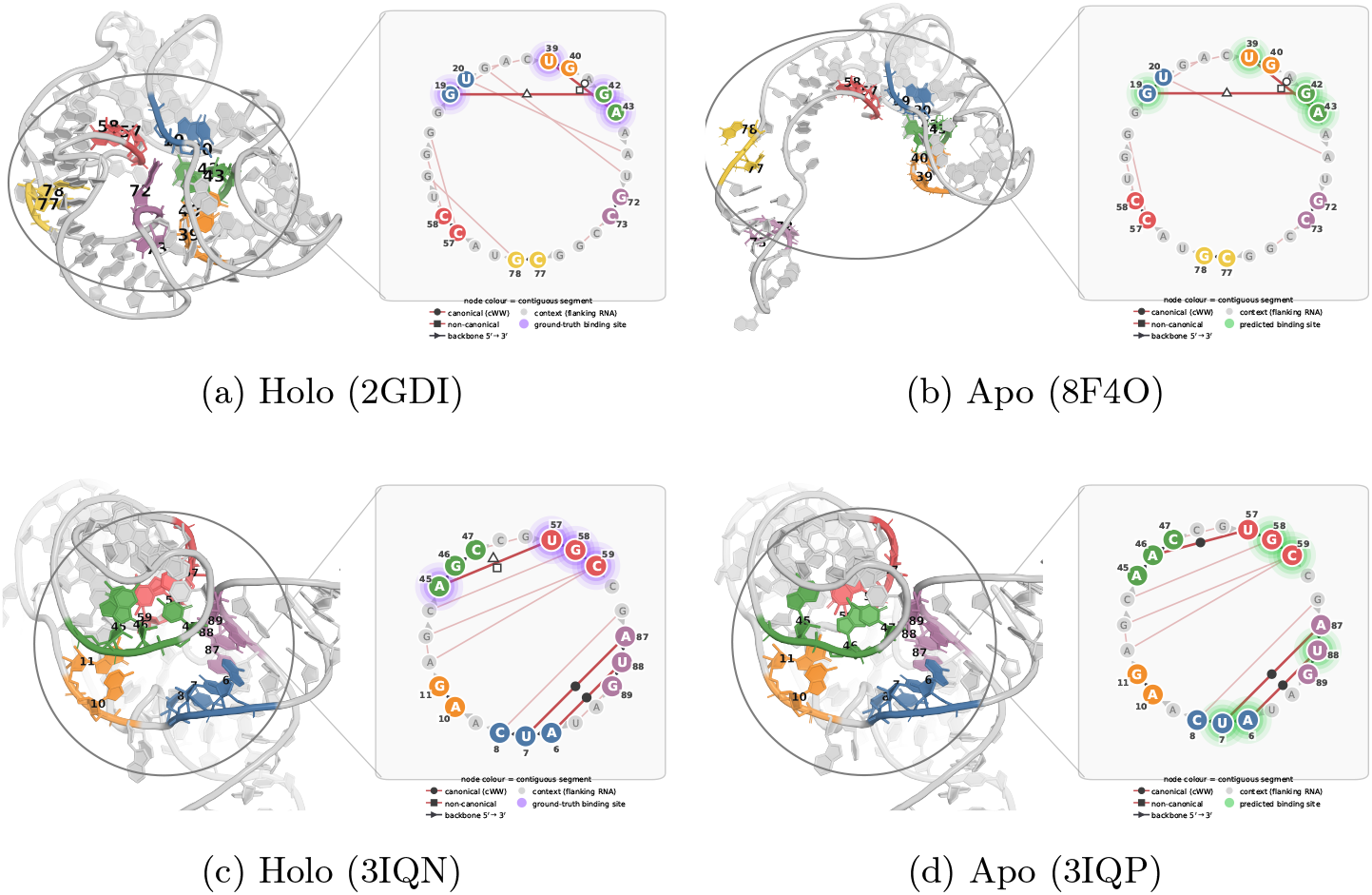
Holo and apo RNA structures with each contiguous segment of the binding-site region drawn in a distinct color, matched between the structure and its 2.5D graph in the magnified panel (residues numbered). Nucleotides are glow-highlighted: on the holo (bound) structures a violet glow marks the ground-truth binding site, and on the apo (unbound) structures a green glow marks the site `Ribo-LENS` predicts from the unbound state. Within each pair the apo structure is superposed onto the holo and both are shown in the same orientation. Top row: 2GDI (holo)/8F4O (apo); bottom row: 3IQN (holo)/3IQP (apo). Comparing across states shows how well the site `Ribo-LENS` recovers from the unbound conformation matches the true bound-state pocket.

**Figure 5:**
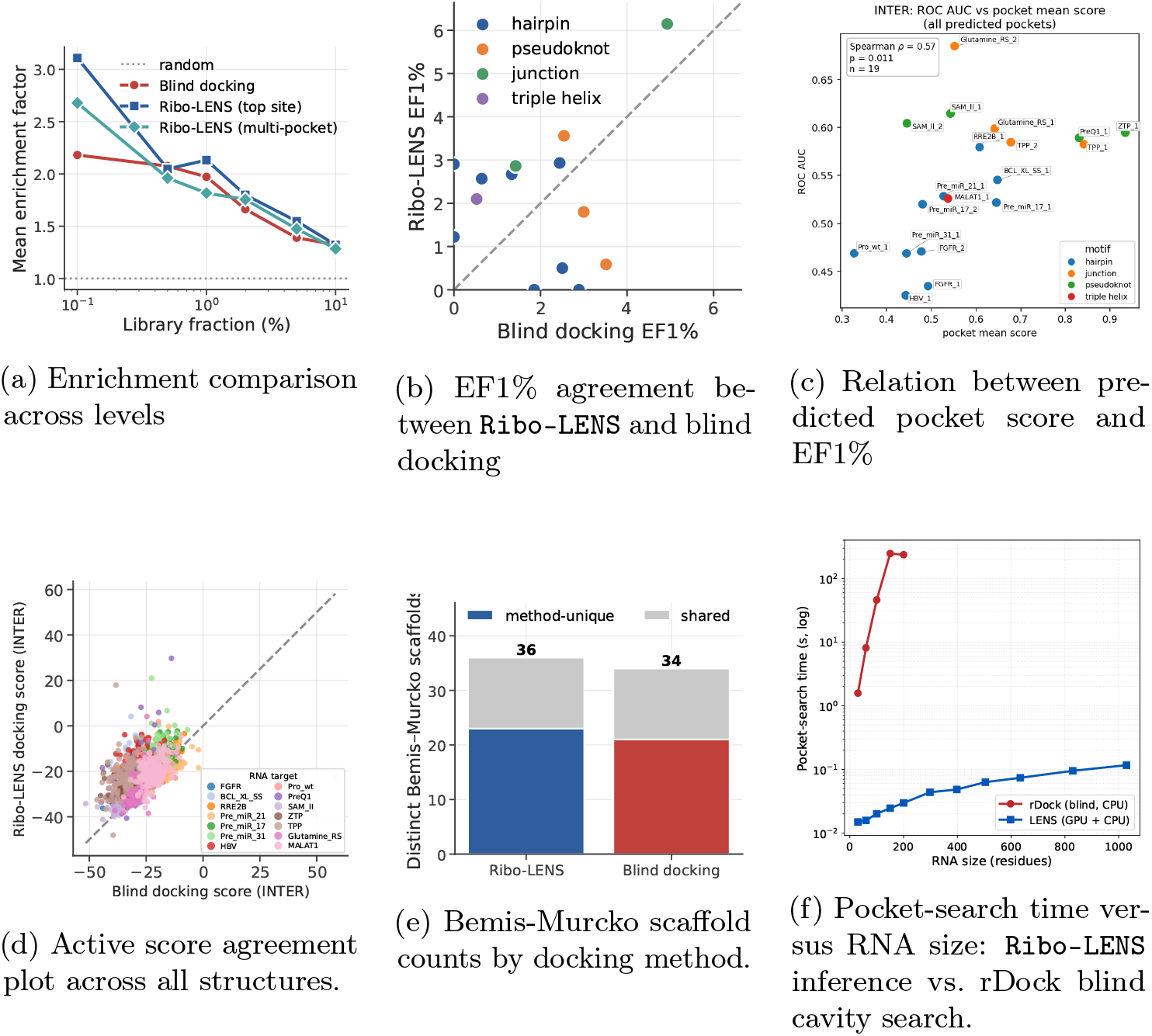
(a) Enrichment factor from blind docking, `Ribo-LENS`, and `Ribo-LENS` multi-pocket (where we use the maximum score across all pockets if multiple are predicted by `Ribo-LENS` on the structure), across fraction of the ranked library. (b) EF1% agreement plot between blind docking and `Ribo-LENS` per structure. (c) Scatter plot between the maximum score assigned to any node in a pocket predicted by `Ribo-LENS` and the enrichment for that structure. There is a statistically significant spearman correlation between the assigned scores and docking enrichment (spearman *ρ* = 0.56, *p* = 0.0087). (d) Active-score agreement across all structures. (e) Bemis–Murcko scaffold counts by docking method. (f) Wall-clock pocket-search time as a function of RNA size (30–1029 nucleotides), on the same machine (AMD Ryzen 9 7950X; `Ribo-LENS` inference additionally using an NVIDIA RTX 5090). `Ribo-LENS` inference (RGCN backbone on GPU followed by edge-pooling) stays in the tens-of-milliseconds range across the whole size range, whereas rDock blind cavity mapping over the entire molecule grows from ∼1.6 s at 30 nt to intractable (*>* 600 s by ∼300 nt). This confirms that blind cavity-search cost scales steeply with RNA size, while pocket-localized `Ribo-LENS` inference does not.

### Ribo-LENS enables a fully sequence-based virtual-screening pipeline

We next test whether `Ribo-LENS` can drive a fully sequence-based virtual-screening pipeline. Using the ROBIN assay ^(5)^ (≈ 25,000 small molecules screened against disease-relevant RNAs), we predict each RNA’s structure from sequence with RhoFold+ ^(18)^, localize sites with `Ribo-LENS` (three pooling steps, no score threshold), and dock the library against a 10 Å sphere around each predicted pocket centroid with rDock ^(19)^. As a baseline we run blind docking, in which rDock detects cavities across the entire structure and the same library is docked into them with no predicted-site prior. We consider the single highest-scoring pocket (*top site*) and an aggregate over all pockets (*multi-pocket*). After removing unfoldable, too-short, or undockable targets (Methods), 14 RNAs (29–84 nt) remain.

### Ribo-LENS is competitive with blind docking’s enrichment at a fraction of the search cost

Both protocols reliably lift active compounds from the library: at the top 1% of ranked compounds the per-structure enrichment factor (EF1%) exceeds the random expectation of 1.0 for both blind docking (*q* = 0.046; one-sided Wilcoxon signed-rank against EF = 1, Benjamini–Hochberg-corrected across all tested methods and levels) and `Ribo-LENS`-guided docking (*q* = 0.037). The two are in fact statistically indistinguishable at every level tested (0.1–5%; Wilcoxon signed-rank, all n.s.), despite `Ribo-LENS` searching only a predicted pocket; per structure (Fig. 5b) `Ribo-LENS` gives the higher EF1% on 9 of 14 targets and the single best value in the panel (EF1%= 6.14 on TPP). The gain is in search cost (Fig. 5f): docking a fixed-radius pocket costs the same regardless of RNA size, so only `Ribo-LENS` inference scales with the molecule, far more cheaply than blind cavity search, which becomes intractable beyond ∼200 nt. Aggregating over multiple pockets (*multi-pocket*) lowered EF at all levels (Fig. 5a), indicating that low-confidence secondary pockets add noise, likely because no score threshold was applied here.

#### Confidence correlates with overall ranking quality

Next, we investigate whether the binding site scores assigned by `Ribo-LENS` correlate with screening performance. Across the 19 pockets docked over the 14 structures, mean pocket score correlates moderately with the whole-ranking quality of the resulting screen (ROC AUC; Spearman *ρ* = 0.57, *p* = 0.011; Fig. 5c), but shows no significant association with top-1% enrichment (EF1%). Pocket confidence is therefore informative for prioritizing targets by expected overall ranking quality, which can be used to triage signal when screening many structures, but is not a reliable predictor of early enrichment specifically.

#### Ribo-LENS and blind docking recover complementary chemotypes

Finally, we asked whether the two protocols surface similar compounds, since chemical diversity, not only binding energetics, is a critical objective in drug discovery^(20)^. Docking scores for true binders are moderately correlated (Spearman *ρ* = 0.63, *p <* 0.01; Fig. 5d) between both tools, yet the chemical matter they surface is largely distinct. Among the actives identified in the top-1% across all structures, the two methods share only 13 of 57 distinct Bemis–Murcko scaffolds (Fig. 5e); for ZTP (ROBIN 31), for instance, the top-1% actives recovered by each protocol are largely distinct (Supp. Fig. 3). `Ribo-LENS`-guided docking identifies 23 unique scaffolds compared to 21 from blind docking. The two approaches therefore recover overlapping but substantially different regions of chemical space: `Ribo-LENS`-guided docking and blind cavity search each surface scaffolds the other misses, making them complementary rather than redundant.

Together, a fully sequence-based pipeline (structure prediction, `Ribo-LENS` localization and pocket-restricted docking) achieves enrichment on par with blind docking while searching only a small predicted region, its confidence scores rank targets by overall screen quality, and the two strategies recover complementary chemotypes, pointing to value in combining them.

### Ribo-LENS localizes binding sites on SARS-CoV-2 RNA elements

A central goal of RNA-targeted drug discovery is to locate druggable pockets within disease-associated structures. As a prospective test of `Ribo-LENS` on therapeutically relevant targets, we examined two highly conserved elements of the SARS-CoV-2 genome: the programmed ribosomal frameshift-element pseudoknot (PK; PDB 6XRZ) and the 5^*′*^-terminal stem-loop 1 (SL1; PDB 9EOW), both proposed as antiviral targets ^(21)^. As throughout, we localize binding sites with three pooling steps at a threshold of 0.5. Neither structure’s sequence is closely related to the training set (maximum identity 0.538 for 6XRZ and 0.613 for 9EOW against the GerNA-Bind training set), so both function as low-homology, held-out targets.

For the frameshift pseudoknot (Fig. 6a), `Ribo-LENS` predicted a single site (A512, G513, U524, G525). Two of these, G513 and G525, are among the four residues with the largest chemical-shift perturbations (CSPs) on ligand binding in NMR titrations ^(21)^, and G525 is independently corroborated by a study with a chemically distinct ligand that localizes the primary site to the S3 stem^(22)^. `Ribo-LENS`’s residues lie within this structured S3 stem rather than a flexible loop, indicating predictions driven by learned interaction geometry rather than a bias toward the most flexible regions.

**Figure 6:**
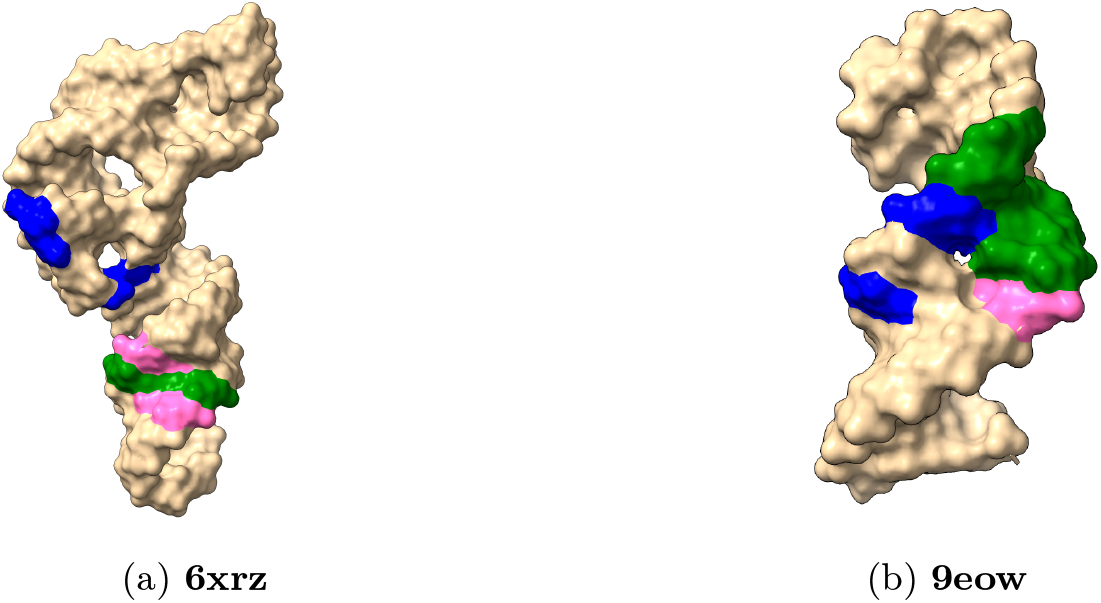
Binding-site prediction comparisons. **(a)** The structure of 6xrz, representing the SARS-CoV-2 PK. **(b)** The structure of 9eow, representing the SARS-CoV-2 SL1. In both panels: blue indicates residues found to have the most significant CSPs when a ligand was introduced, green indicates residues predicted to be the binding site by our model, and pink indicates residues that had the highest CSP and were successfully predicted as a binding site by our model.

For SL1 (Fig. 6b), which presents an internal loop and a hairpin loop, `Ribo-LENS` identified the internal loop, predicting residues 25–28 and directly recovering C28, one of the three residues (U11, U13, C28) with the largest CSPs^(21)^. Further pooling expands around the same internal loop rather than migrating elsewhere, indicating the model concentrates on a single coherent motif.

Together, these held-out, low-homology cases show `Ribo-LENS` recovering experimentally implicated binding regions; its agreement with independent NMR data across two structurally distinct SARS-CoV-2 elements supports its use for nominating candidate sites in RNA-targeted drug-discovery campaigns.

## Discussion

We introduced `Ribo-LENS`, a method for predicting small-molecule binding sites in RNA. Two features set it apart. First, it works entirely from the 2.5D base-pairing graph, a coarse-grained representation that discards atomic coordinates yet retains the interaction geometry that defines a pocket, making it robust to conformational change and usable wherever only low-resolution or predicted structure is available. Second, it reasons over connected substructures rather than individual residues: by contracting edges and pooling the nucleotides they join, it forms and scores representations of connected *groups* directly, so each step asks whether a connected set of residues constitutes a binding site rather than scoring residues in isolation and clustering them post hoc. Of the two, we regard the substructure-level formulation as the more general methodological contribution, a principle that should transfer to other problems where the object of prediction is a connected motif rather than a set of independent sites.

Strikingly, and despite drawing on far less information, `Ribo-LENS` outperforms methods that depend on accurate 3D structure (RNAsite), fine-tuned language models (GerNA-Bind), and all-atom co-folding models (AlphaFold3, Chai-1) on a held-out benchmark, and matches an explicit all-atom cavity search in end-to-end screening. This resilience is structural in origin: secondary structure forms early in RNA folding^(23)^ and the base-pair network is comparatively stable across the apo-to-holo transition, so `Ribo-LENS`’s predictions track the base-pairing graph rather than 3D geometry, holding under even large-RMSD rearrangements and shifting only when binding rewires the pocket itself. Encouragingly, the signal it learns from crystallographic data alone is echoed by three independent lines of evidence: structure-based docking, the *in vitro* ROBIN binding assay, and NMR chemical-shift perturbations. This indicates it captures genuine binding determinants.

Our approach has limitations. `Ribo-LENS` localizes pockets but does not itself model binding-site flexibility or the induced fit that accompanies ligand binding, and rearrangements that reshape the apo base-pairing network will degrade it; learning over an ensemble of plausible network reconfigurations is a natural next step. Because it consumes structure rather than producing it, `Ribo-LENS` also inherits the errors of whatever structure source it is paired with, albeit tolerantly given its coarse-grained input. A further priority is to validate the approach at scale against experimentally confirmed binders.

`Ribo-LENS` is not a replacement for 3D-based methods or co-evolutionary language models; its value is in operating where they cannot, on the majority of RNAs that lack a high-resolution structure or a strong homology signal, as a rapid first pass that triages targets for closer 3D scrutiny. By publicly releasing our datasets, source code and model weights, we hope to catalyze a community effort in this direction. Notably, `Ribo-LENS` has the distinct advantage of enabling structure-based virtual screening from only low-resolution structural information such as base-pairing interactions: predicted structures, secondary-structure models from tools such as SPOT-RNA^(24)^ and ViennaRNA ^(25)^, or chemical-probing data. Paired with rapidly emerging RNA-centric design technologies and with all-atom models such as AlphaFold3 ^(12)^ that now support nucleic acids, `Ribo-LENS` can help pave the way for the next generation of RNA drug discovery. In light of the daunting number of potential RNA targets, screening from coarse structural signal alone could prove a major asset for mining entire transcriptomes and fully embracing the era of RNA therapeutics.

## Methods

### Datasets

We evaluate `Ribo-LENS` on three datasets that probe complementary aspects of binding-site prediction: held-out structural generalization (GerNA-Bind), robustness to conformational state (Holo–Apo Pairs), and downstream virtual-screening utility (ROBIN). The construction pipeline for the training data (retrieval from the PDB, distance-based labeling, 2.5D-graph conversion, and redundancy-aware structure-based splitting) is summarized in Fig. 1c.

#### GerNA-Bind

We use the GerNA-Bind dataset^(15)^, derived from RNA–ligand complexes in the HARIBOSS database^(26)^. Each entry is a single RNA chain, and the dataset is partitioned into training, validation, and test splits by a temporal cutoff: complexes deposited before 2020 constitute the training set, and those from 2021 onward the validation and test sets, with 2020 held out as a buffer between splits. Ground-truth labels are released only for the test split; for training and validation we therefore label as positive any nucleotide with at least one atom within 4 Å of a ligand atom, computed in ChimeraX^(27)^. The test set retains the labels of the GerNA-Bind benchmark which use contact labels between the ligand and RNA.

#### Holo-Apo Pairs

To assess robustness to conformational state, we use the holo/apo benchmark introduced with fpocketR^(4)^, comprising ten RNAs each resolved in both a ligand-bound (holo) and ligand-free (apo) form. We exclude one pair whose structure (>2,900 nucleotides) exceeds the largest training example (209 nucleotides) by more than an order of magnitude, leaving nine pairs. This benchmark tests whether the model can generalize to a more realistic setting, since at deployment we are usually given an apo state, whereas the training data naturally consists of holo structures.

#### ROBIN

To evaluate downstream screening utility, we use the ROBIN assay^(5)^, an experimental library of ≈ 25000 small molecules screened against nucleic-acid (RNA and DNA) targets. We use it to compare structure-based virtual screening when docking is restricted to the binding site predicted by `Ribo-LENS` against blind docking over the entire structure, quantifying performance by the enrichment factor among the top 1% of docking scores (EF1%).

### RNA Graph Representation

We use RNAGlib to represent each RNA as a 2.5D graph *G* = (*V, E*). Nodes correspond to nucleotides and carry a one-hot encoding of nucleotide identity concatenated with a 640-dimensional residue embedding from the RNA-FM language model^(16)^, which was pre-trained on a large corpus of RNA sequences and provides a strong evolutionary signal. Edges encode chemical interactions following the Leontis-Westhof classification^(7)^, a twelve-family geometric taxonomy of base pairs defined by the interacting edges of each base (Watson–Crick, Hoogsteen, and sugar) together with the cis/trans orientation of the glycosidic bonds. We augment these base-pairing edges with backbone (5^*′*^ → 3^*′*^) connectivity and represent all interactions as *directed* edges, which captures sequence orientation and the intrinsic asymmetry of non-canonical pairs. This yields a relation set ℛ of 20 distinct edge types. This graph-native, geometry-aware representation is well matched to relational message passing, which we exploit below.

### Model Architecture

#### Relational Graph Convolutional Network (RGCN)

We process the input graph with Relational Graph Convolutional Network (RGCN) layers ^(17)^, which learn a separate transformation per edge type and are therefore well suited to our typed RNA graph. Let ℛ be the set of relations and *N*_*r*_(*i*) the neighbors of node *i* under relation *r*. A layer *l* is parameterized by |ℛ| + 1 matrices 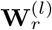, where index 0 denotes the self-loop. With 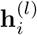 the representation of node *i* at layer *l*,

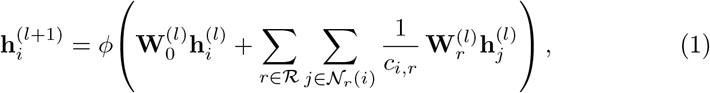

where *c*_*i,r*_ = | *N* _*r*_(*i*) | is a per-relation normalization constant and *ϕ* is a non-linear activation. We stack 4 RGCN layers. Each input node feature (dimension 644) is first mapped to dimension 256 by a linear projection. The final layer produces, for every node *i*, an embedding **z**_*i*_ and a scalar binding score 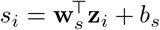.

#### Edge Scoring and Contraction

Our pooling module follows the edge-contraction structure of EdgePool ^(14)^ but, unlike the original end-to-end gated formulation, learns edge scores under direct supervision (see Training). For each node we form the pooling feature **n**_*i*_ = **z**_*i*_ ∥ (*s*_*i*_ −*s*_*μ*_) by concatenating its RGCN embedding and binding score minus the mean of the scores (*s*_*μ*_). The score of edge *e*_*ij*_ is

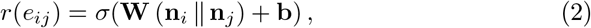

with *σ* the logistic sigmoid, so *r*(*e*_*ij*_) ∈ (0, 1). When an edge is contracted, its two endpoints are merged into a single supernode whose feature is the size-weighted average

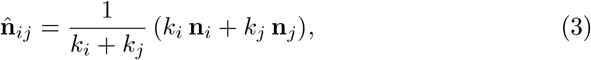

where *k*_*i*_ is the number of original nucleotides represented by node *i*. As a node absorbs more nucleotides over successive contractions, its feature is weighted accordingly.

### Binding-Site Prediction with Ribo-LENS

Existing RNA binding-site predictors typically operate at the level of individual nucleotides, assigning each residue an independent binding probability. Small-molecule pockets in RNA, however, are not collections of independently druggable nucleotides; they are defined by a local network of interactions among residues. A binding site is therefore better viewed as a connected subgraph motif in the interaction network than as a set of high-scoring nodes.

`Ribo-LENS` captures this by coupling the RGCN with a learned graph-coarsening procedure that reasons over interactions rather than residues. The RGCN is evaluated *once* to produce the node features *{***n**_*i*_ *}*; the pooling module then iterates. At each step it scores every current edge with Equation 2 and contracts the single highest-scoring edge whose score exceeds a threshold *P* = 0.5, merging its endpoints via Equation 3; the coarsened graph is then re-scored and the process repeats. Crucially, contraction recruits edges, not isolated nodes, so it grows outward from a high-confidence core interaction and progressively condenses contiguous subregions into coherent clusters. The procedure terminates when no edge exceeds *P* or after *T* steps. Every resulting supernode that spans two or more nucleotides is reported as a predicted binding region, and a nucleotide is labeled positive exactly when it has been absorbed into such a supernode.

This formulation offers two advantages over node-level classification. First, it enforces spatial and topological coherence: predicted residues must be connected through interactions rather than scattered across the structure. Second, the iterative contraction provides a natural mechanism for pocket-boundary delineation, growing a prediction from a high-confidence core until the local binding signal is exhausted.

### Training

We train `Ribo-LENS` in two stages: the RGCN is trained to convergence and frozen, after which the pooling module is trained on top of it.

#### Stage 1: RGCN

The RGCN receives the 2.5D graph (RNA Graph Representation) and is trained to predict per-nucleotide binding labels with a binary cross-entropy (BCE) logits loss on the node scores *s*_*i*_. We select the checkpoint with the highest validation AUPRC rather than the lowest validation loss, for two reasons. First, with strong class imbalance (*<* 20% of nodes are positive), BCE can be reduced by improving on the prevalent negatives while degrading the positive class we care about. Second, the downstream pooling stage performs greedy contraction over RGCN scores and embeddings, so positive-class ranking is the quantity that governs downstream behavior. We stop when validation AUPRC has not improved for 10 epochs and freeze the RGCN.

##### Algorithm 1

Binding-site prediction via `Ribo-LENS` pooling (inference)

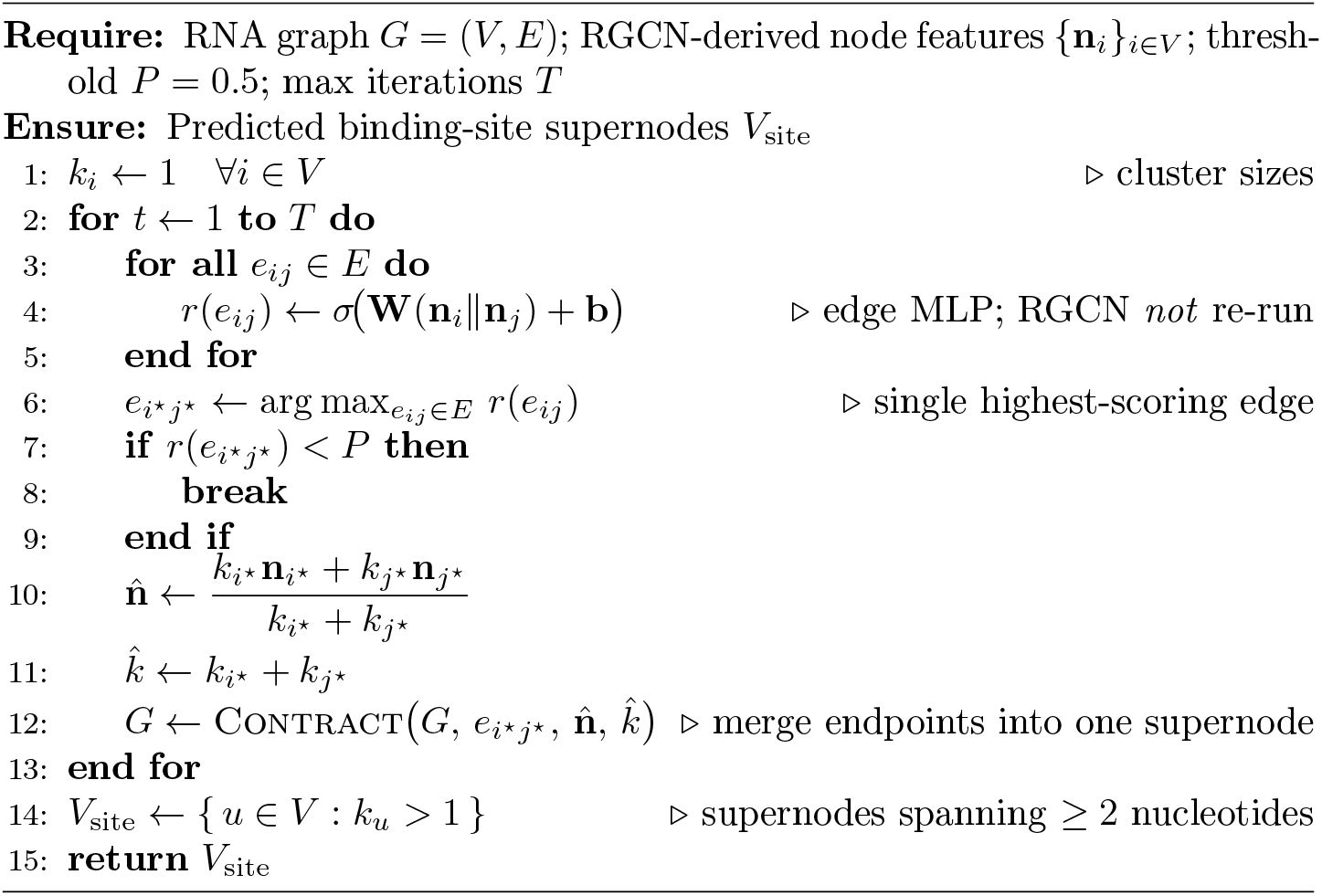

#### Stage 2: Pooling

With the RGCN frozen, its node features are passed to the pooling MLP, which is trained with BCE against the soft edge target

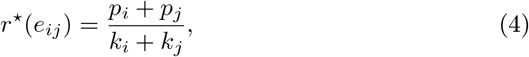

where *p*_*i*_ is the number of binding-site nucleotides and *k*_*i*_ the total number of nucleotides represented by node *i*; thus *r*^⋆^ is the fraction of binding-site nucleotides spanned by the contracted pair. For each training batch we perform three rounds of pooling so the model also sees coarsened graphs. Within a round we score all edges and contract a random number (between one and 10% of |*E*|) of the highest-scoring edges, skipping any edge that shares a node with an edge already selected in that round. This stochastic, multi-edge contraction is a cheap and higher-variance proxy for the single-edge greedy contraction used at inference (Algorithm 1) and exposes the MLP to a wider range of coarsened states. We again select on validation AUPRC, treating an edge as a positive when *r*^⋆^(*e*_*ij*_) ≥ 0.5.

### Evaluation metrics

We evaluate binding-site prediction at the nucleotide level using Matthews correlation coefficient (MCC), precision, recall and F1, treating each nucleotide as binding or non-binding. We report MCC as the primary metric because it is well suited to the strong class imbalance in this task (binding nucleotides are a small minority of each structure). Where a threshold-independent view is informative, we additionally report the area under the precision–recall curve (AUPRC), which characterizes performance across the positive class over all decision thresholds and, like MCC, is appropriate under class imbalance.

We also use the Spearman rank correlation coefficient *ρ*, which measures the strength of a monotonic relationship between two variables without assuming linearity and report the associated *p*-value.

For the virtual-screening experiments we quantify early recognition with the enrichment factor (EF). For a given target and a top fraction *x* of the docking-ranked library, the enrichment factor is

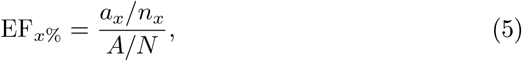

where *n*_*x*_ is the number of compounds in the top *x*% of the ranking, *a*_*x*_ the number of experimentally confirmed actives among them, and *A/N* the fraction of actives in the full library of size *N* .

## Data Availability

The datasets and model checkpoints can be downloaded here

## Code Availability

The code for training and testing the models is available at Ribo-LENS

## Author Contributions

D.N. implemented the model, conducted the experiments, contributed to algorithm design, and contributed to writing and figures. J.W. conceptualized the research, contributed to writing, acquired funding, and supervised the project.

C.O. conceptualized the research, led the model design, contributed to writing, acquired funding, and supervised the project.

## Supplementary Information

**Supplementary Figure 1:**
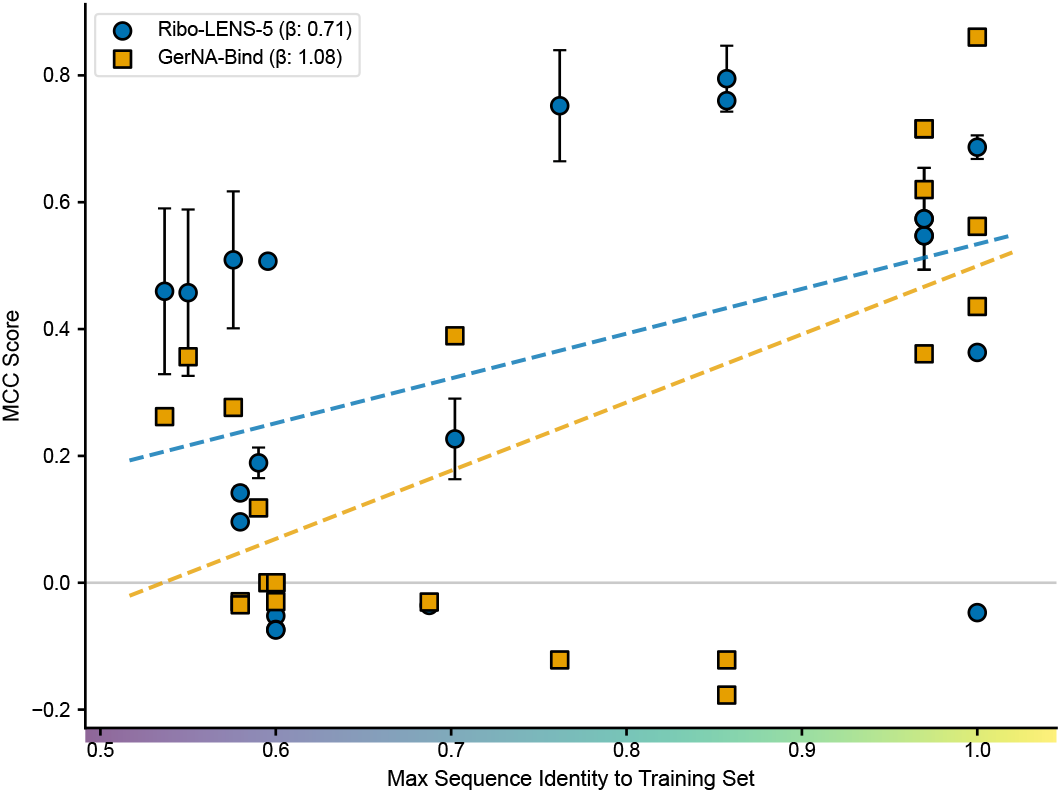
Per structure MCC vs Homology scatter between `Ribo-LENS` and GerNA-Bind at 5 pooling steps

**Supplementary Figure 2:**
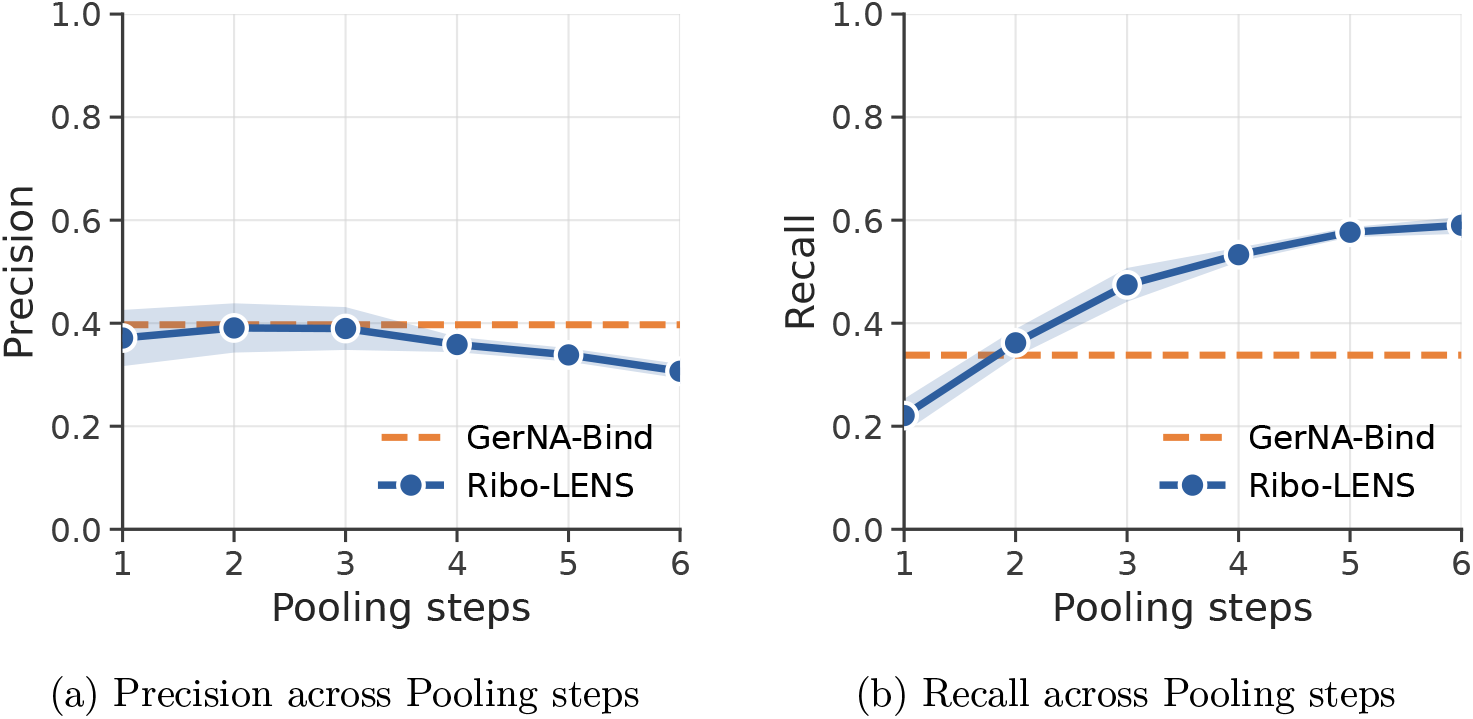
Precision and recall on the GerNA-Bind test set as a function of pooling steps. **(a)** Precision and **(b)** recall for `Ribo-LENS` (blue; shaded band, standard deviation across 9 seeds) and GerNA-Bind (orange dashed line). Precision tracks GerNA-Bind through the first three pooling steps and then erodes, while recall increases monotonically. The corresponding MCC sweep is shown in the main text (Fig. 2c).

**Supplementary Figure 3:**
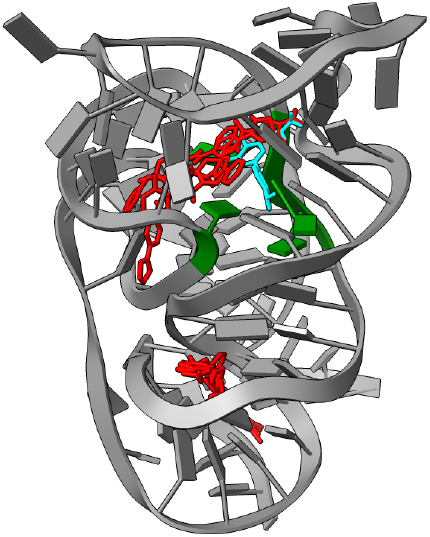
The 3D structure of ZTP (ROBIN 31) used for docking. Ligands in red are actives in the top 1% of scores from blind docking; ligands in cyan are actives in the top 1% of scores from `Ribo-LENS`-guided docking; residues colored in green are the binding site identified by `Ribo-LENS`.

**Supplementary Figure 4:**
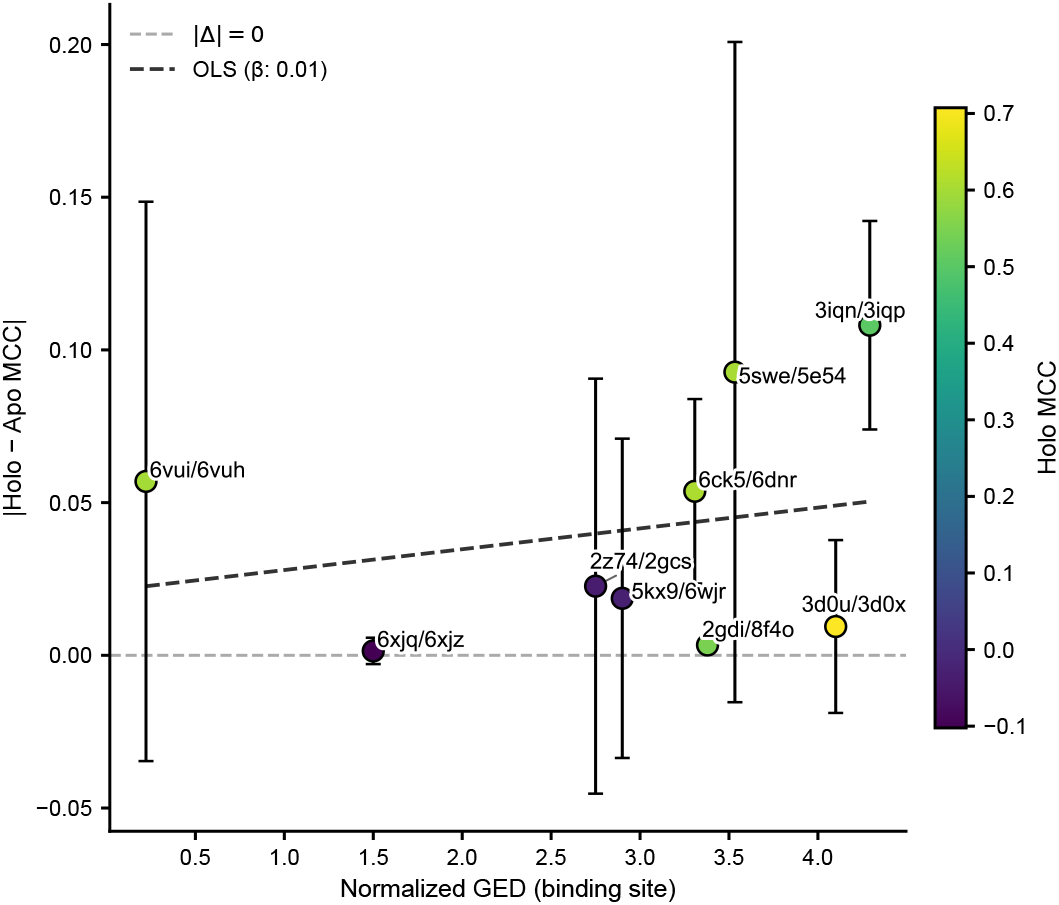
Holo–Apo pair absolute MCC delta vs 1hop normalized GED scatter

**Supplementary Figure 5:**
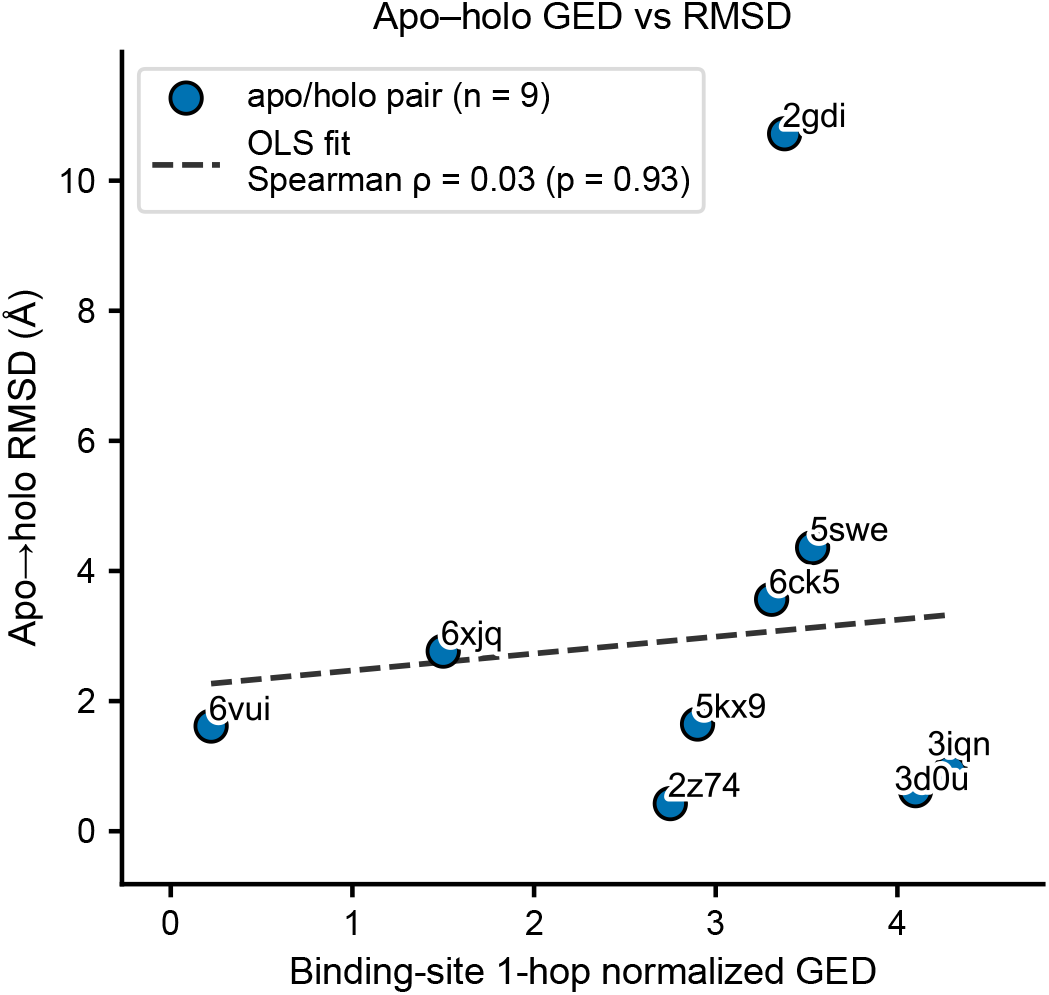
Holo–Apo 1-hop GED correlation with RMSD

